# Single Cell RNA Sequencing Reveals Heterogeneity of Human MSC Chondrogenesis: Lasso Regularized Logistic Regression to Identify Gene and Regulatory Signatures

**DOI:** 10.1101/854406

**Authors:** Nguyen P.T. Huynh, Natalie H. Kelly, Dakota B. Katz, Minh Pham, Farshid Guilak

## Abstract

Bone marrow-derived mesenchymal stem cells (MSCs) exhibit the potential to undergo chondrogenesis *in vitro*, forming *de novo* tissues with a cartilage-like extracellular matrix that is rich in glycosaminoglycan and collagen type II. However, it is now apparent that MSCs comprise an inhomogeneous population of cells, and the fate of individual subpopulations during this differentiation process is not well understood. We analyzed the trajectory of MSC differentiation during chondrogenesis using single cell RNA sequencing (scRNA-seq). Using a machine learning technique – lasso regularized logistic regression – we showed that multiple subpopulations of cells existed at all stages during MSC chondrogenesis and were better-defined by transcription factor activity rather than gene expression. Trajectory analysis indicated that subpopulations of MSCs were not intrinsically specified or restricted, but instead remained multipotent and could differentiate into three main cell types: cartilage, hypertrophic cartilage, and bone. Lasso regularized logistic regression showed several advances in scRNA-seq analysis, namely identification of a small number of highly influential genes or transcription factors for downstream validation, and cell type classification with high accuracy. Additionally, we showed that MSC differentiation trajectory may exhibit donor to donor variation, although key influential pathways were comparable between donors. Our data provide an important resource to study gene expression and to deconstruct gene regulatory networks in MSC differentiation.

## Introduction

Articular cartilage is an avascular and aneural connective tissue that exhibits little to no intrinsic capability for repair (1). Therefore, cartilage defects caused by injury or progressive degeneration in the case of diseases such as osteoarthritis often result in long-term pain and disability (2). While there are currently no disease-modifying treatments available for osteoarthritis, significant efforts have been ongoing in the field of regenerative medicine to create tissue engineered constructs that can mimic the mechanical and biological characteristics of native articular cartilage.

Among many potential cell sources for cartilage regeneration, bone marrow derived mesenchymal stem cells (MSCs) provide an attractive system due to their accessibility and their capacity for *in vitro* expansion (3, 4). However, several studies have found evidence that MSCs grown under chondrogenic conditions (e.g., pellet aggregate) tend to follow the developmental process of hypertrophic differentiation and endochondral ossification, where resultant tissue exhibits high content of collagen type I and type X (5, 6). Indeed, transcriptomic profiling of MSC differentiation has indicated that canonical markers for chondrogenesis, hypertrophic differentiation, or osteogenesis (COL2A1, COL10A1, COL1A1) increased simultaneously throughout this process (7). The ability to separate gene regulatory networks (GRN) underlying these differentiation routes is highly valuable, as a comprehensive understanding of decision points may enable more effective protocols in cartilage tissue engineering to prevent unwanted phenotypes (e.g., osteoblastic or hypertrophic).

Decoupling the signaling pathways that induce a desired phenotype is central to the understanding of how alternative protocols could be enhanced. Attempts to decipher the GRN from bulk RNA-sequencing have failed to untangle pathways involved in chondrogenesis from hypertrophy (7), likely due to the fact that MSCs comprise multiple cell types and bulk RNA-seq measures the average expression of all cell types present. On the contrary, state-of-the-art single cell RNA sequencing (scRNA-seq) provides a means of determining cellular identity and complexity at single cell resolution, where profiles of gene expression and transcriptional programs of each cell type can be determined. Moreover, data from scRNA-seq can also be used to construct a proposed differentiation trajectory based on decision points at which multipotent cells progress towards one identity versus the other.

One challenging task in scRNA-seq analysis is the identification of key genetic patterns for the classification of cell identities from high dimensional gene expression profiles. Compared to traditional strategies of microarray or bulk RNA-seq, scRNA-seq makes it possible to apply prediction methods such as logistic regression to solve this classification difficulty by overcoming the hurdle of limited sample size. However, logistic regression requires a thorough process of model selection to pinpoint the few genes that discriminate one cell type from another, which is usually computationally challenging. In addition, the number of features is very high, making logistic regression classifiers unreliable. To tackle this problem, we instead examined the use of lasso regularized logistic regression (LRLR) for feature selection and model building. LRLR is a machine learning technique for high dimensional data that combines the discriminative power of logistic regression and the ability to perform variable selection from regularization methods. Not only does LRLR provides classifiers to distinguish between cell types, it also automatically identifies key influential genes. LRLR results are usually parsimonious in that they contain very few important features, making it easy to focus on the most influential targets.

In this study, we utilized scRNA-seq to describe the GRNs and cellular trajectories of MSCs during chondrogenic differentiation. More importantly, we applied LRLR to identify novel markers that were specific to each differentiated cell type, as well as to pinpoint distinct transcriptional programs governing these processes. LRLR results were then applied to determine influential signaling pathways that could enhance the outcome of engineered cartilage.

## Results

### Transcriptomic programs are distinguished from gene expression

MSCs were cultured in 3D pellet aggregates in chondrogenic medium and samples were collected at 4 different time points during chondrogenesis (day 0, day 1, day 7, and day 14). We captured a total of ~35,000 cells across four time points, with 2324 ± 654 median reads per cell. Initial cluster analysis identified 3-4 subpopulations of cells per sample based on gene expression. However, it had been previously suggested that cell state characterization on the level of regulatory network may be more advantageous, since this method could overcome dropouts and technical variation (8). Thus, we further characterized cellular architecture based on transcription factor activities (TFA) and identified 3-4 subpopulations within each timepoint (Figure S1A). We set out to elucidate whether subpopulations during MSC chondrogenesis may be most represented by gene expression (GE) or by transcription factor activity (TFA).

#### Day 0

Cells at day 0 expressed canonical mesenchymal markers, such as CD44, ENG (CD105), THY1 (CD90), and NT5E (CD73), in concordance with previous flow cytometry results from our group (Figure S1B) (7). Typical markers of hematopoietic lineages (CD34, CD19, CD45, CD11b) were not expressed in our data set. Cells were clustered into three subpopulations based on gene expression (GE1 – GE3), while there appeared to be four distinguished regulatory identities (TFA1-TFA4) (Figure S1A and 1A). Specifically, GE1 was further divided into TFA3 and TFA4, suggesting while there may be similarity in gene expression profiles, different cells may still exhibit very distinguishable transcriptional programs. Furthermore, we showed that such transcription factor activity, defined by the co-expression of all its downstream targets, was distinguished from the expression of the transcription factor itself (Figure 1B).

**Figure 1:**
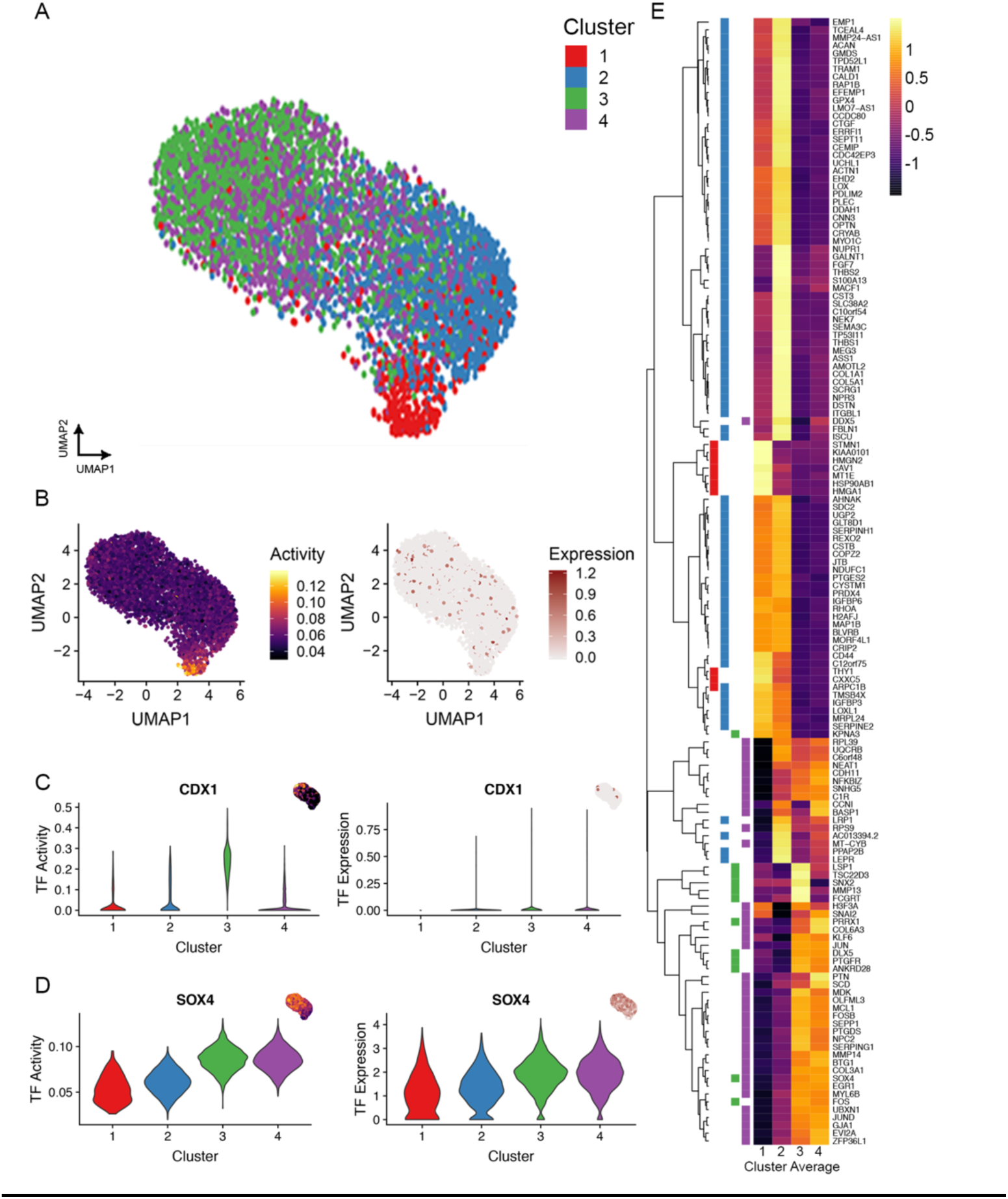
Gene and Regulon Signature of Day 0 Subpopulations. (A) Cellular clustering based on transcription factor (TF) activity. (B) Transcription factor activity was distinguished from transcription factor expression. (C-D) Violin plots showing the top transcription factor for TFA3 and TFA4 and their activities versus expressions. (E) Heatmap depicting gene signatures for each cluster, color-mapped by gene average expression per cluster.

Next, we utilized lasso regularized logistic regression (LRLR) to decipher gene markers characteristic of each TFA cluster. Compared to the popular method of the Wilcoxon rank sum test, we found that LRLR identified markers that were more specific (low overlapping rate between cluster markers) (Figure S2A), and generally fewer number of genes (Figure S2B), whose cluster expressions were well-defined and grouped into four cell states by hierarchical clustering (Figure 1E compared to Figure S2C). Among the MSC canonical surface markers, THY1 and CD44 were identified by LRLR for TFA1 and TFA2 respectively (Figure 1E).

However, their coefficients were relatively lower compared to other cluster markers (SI 1). Interestingly, collagen molecules and matrix metalloprotease molecules were also differentially expressed among subpopulations: COL1A1 and COL5A1 in TFA2; MMP13 in TFA3; COL3A1, COL6A3 and MMP14 in TFA4 (Figure 1E). To further elucidate the regulatory networks involved in these differences, we used LRLR to pinpoint the transcription factors whose regulons (a network of a transcriptional program and its downstream targets) would highly influence each cluster. For example, we found that CDX1 was an influential transcription factor for TFA3, and regulated MMP13 expression (Figure 1C). CDX1 had previously been implicated in skeletal development, acting to relay retinoic acid signals (9, 10); and MMP13 is required for normal development of growth plate cartilage (11). Meanwhile, SOX4 was heavily weighted for TFA4, and regulated MMP14 and COL3A1 expression (Figure 1D). While Cdx1 has previously been shown to be expressed in the forelimb bud region, Sox4 had been shown to exhibit high levels in mesenchymal tissues in the mouse (10, 12). Thus, we speculate that TFA clusters may represent MSCs at different time points in development. In particular, TFA4 may be composed of progenitors at an earlier, more naïve state and TFA3 may represent multipotent cells arisen later. While clustering based on gene expression identified cells from TFA3 and TFA4 as one cluster, these two subpopulations were indeed distinct, demonstrated by the identification of two separate factors that resulted in differential gene expressions between TFA3 and TFA4. Thus, our results highlighted and advocated for the utilization of GRN inference to guide subpopulation discovery.

#### Day 1 and Day 7

While we viewed day 1 and day 7 as transitional states and utilized these samples below for trajectory analysis, we also reported heavily weighted TFs and gene markers of these time points in SI 1.

#### Day 14

We identified three clusters based on gene expression and four clusters based on transcription factor activities for cells on day 14 (Figure 2A and S1A), indicating that MSC chondrogenesis is a heterogeneous process that resulted in many differentiated cell states. In order to elucidate the resulting cell types, we first attempted to classify cells by key marker genes. Cells that expressed LUM or APOE were labeled “stromal” or “adipose”, respectively; cells that exhibited ACAN and MATN4 while maintaining low levels of both COL10A1 and COL1A1, “cartilage”; and cells that highly expressed COL2A1, COL10A1, and COL1A1, “hypertrophic” (short for “hypertrophic cartilage”) (Figure 2A). Our second approach was to predict cell type based on the mouse joint atlas (13), by comparing MSC profiles to cells isolated from murine developing joints (14) (Figure S1C). Both methods led us to identify TFA1 as the hypertrophic cluster, and TFA3 as the cartilage cluster. Cells constituting TFA4 bore resemblance to both immature and early osteoblasts, also expressing both LUM and COL1A2. Thus, we termed TFA4 the stromal/early bone cluster. Once again, network inference successfully separated distinct biological cell states that would otherwise have been missed by solely investigating gene expression.

**Figure 2:**
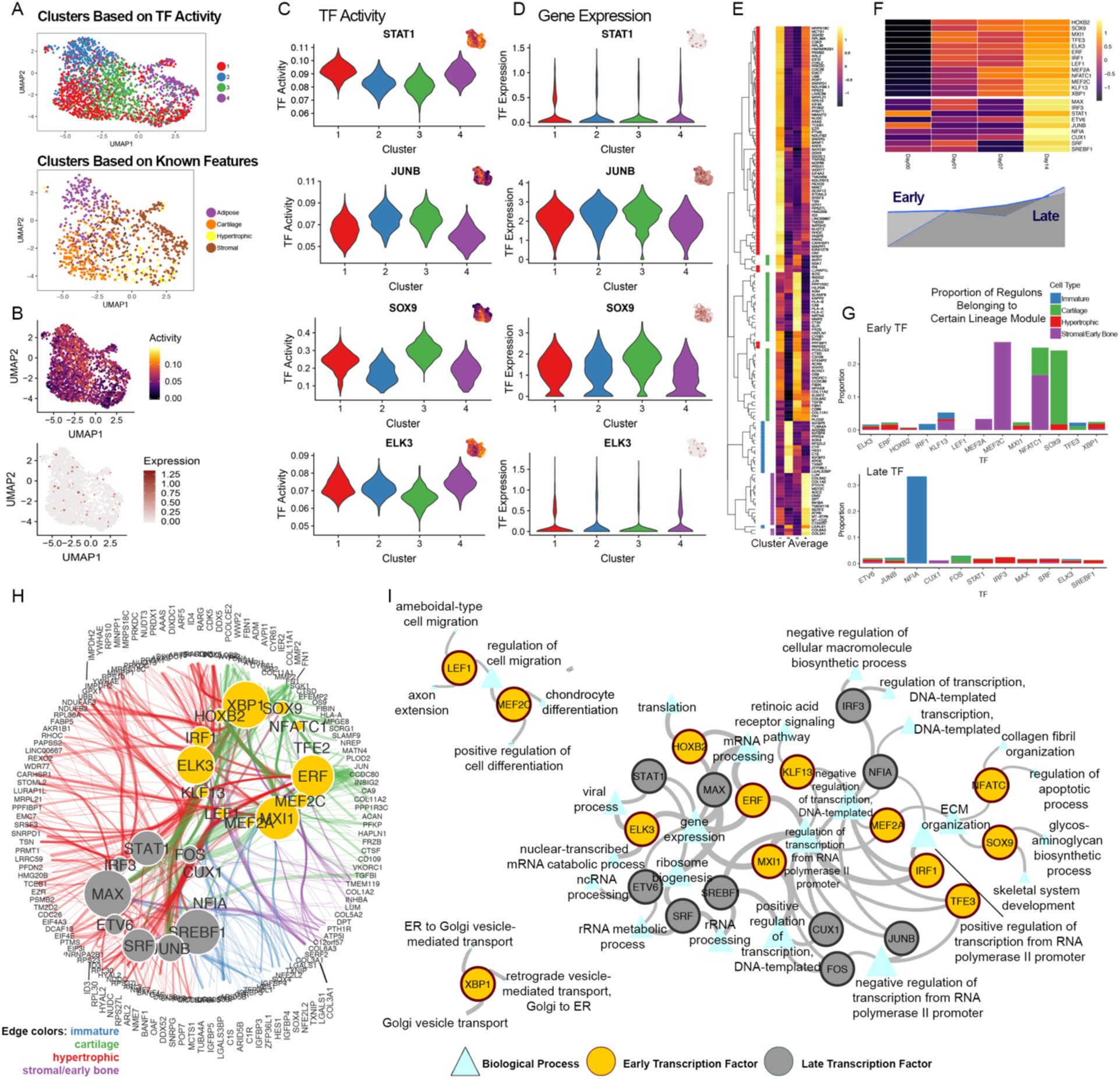
Gene and Regulon Signature of Day 14 Subpopulations. (A) Cellular clustering based on transcription factor activity, or based on supervised classification using known tissue features. Dotted circle in lower panel depicts area of TFA3, indicating this sub-cluster is composed of early stromal or cartilage-like cells. (B) Transcription factor activity was distinguished from transcription factor expression. (C-D) Top transcription factor for each cluster as depicted by their activities and expressions. (E) Heatmap depicting gene signatures for each cluster, color-mapped by gene average expression per cluster. (F) Heatmap depicting early and late regulon signatures for each cluster and their activity. (G) Proportion of early and late regulon targets that belong to immature, cartilage, hypertrophic or early bone gene signatures. (H) A network of early and late transcription factors with their respective targets. Network edges were colored by lineage-related gene signature. Node size signified the number of connections each transcription factor exhibited. (I) A network of early and late transcription factors and the inferred biological processes they controlled. Edge width increased with p-value significance. Triangle node size signified the number of targets in each biological process.

Similar to day 0, we showed that transcription factor activity was distinguished from its expression (Figure 2B). Among the heavily weighted TFs for each cluster (Figure 2C-D), JUNB, an effector for glucocorticoid response, appeared to control the adipose cell fate. It had been previously established that glucocorticoids were critical for chondrogenic differentiation of MSCs, by inducing the expression of many cartilage matrix components (15, 16). Therefore, the presence of JUNB activity suggested that TFA2 was likely to represent an immature state instead of an adipogenic lineage. In addition, SOX9 is a well-established transcription factor in chondrocyte differentiation, and was indeed identified to be directing the cartilage cluster (TFA3). Interferon signaling by IRF3 and STAT1 seemed to be driving the hypertrophic identity (TFA1), and ELK3 led early bone state (TFA4), although their roles in MSC differentiation are not fully understood. By LRLR, we also pinpointed cluster-specific genes and termed these lineage modules, since they represented diverse specification states during MSC differentiation. Interestingly, while a canonical marker for chondrocytes, COL2A1 was not among our lineage module, as its expression was similar between hypertrophic (TFA1) and cartilage (TFA3) subpopulations (Figure S1D). On the other hand, we found many genes related to collagen molecules and extracellular matrix heavily weighted for TFA3, such as ACAN, MATN4, COL11A1, COL11A2, and COL6A1 (Figure 2E). All together, these constituted gene signatures that were specific for immature, cartilage, hypertrophic cartilage, and early bone.

Next, we attempted to reconstruct the gene regulatory networks for each lineage. During MSC differentiation, there appeared to be two waves of transcription factors: the early TFs whose activities increased at day 1 and stayed elevated during differentiation, and the late TFs whose activities did not surge until after day 7 (Figure 2F). As each TF forms a regulatory network (i.e., regulon) with its downstream targets, the composition of these regulons could indicate whether a TF was lineage-specific or not (Figure 2G). By this means, we found that even early on during differentiation, there had been TFs specified in regulating solely the early bone lineage (MEF2A and MEF2C). Other early TFs appeared to be multipotent, since their regulated targets belonged to more than one lineage. Of note, SOX9 – a canonical TF for chondrocyte differentiation – was bipotential between cartilage and hypertrophic, although its hypertrophic targets appeared to be fewer compared to its cartilage targets. On the contrary, late TFs were more specialized, with most of them only regulating targets in one lineage (NFIA: immature, CUX1: early bone, FOS: cartilage, STAT1, IRF3, MAX: hypertrophic). Utilizing regulon information, we built a gene regulatory network of early and late TFs, highlighting the intricate cross-talk and regulation during MSC differentiation (Figure 2H).

We also characterized early and late regulons using gene ontology analysis on each TF’s targets and presented the three most significant biological processes (Figure 2I). Here, the network indicated that highly connected nodes were pathways involved in transcription and translation, signifying processes critical for the synthesis and *de novo* assembly of the extracellular matrix. Other pathways corroborated previous studies on cartilage development, especially SOX9 in glycosaminoglycan synthesis and skeletal system development (17–19), or NFATC1 in collagen fibril organization (20). These results also aligned with the composition of SOX9 and NFATC1 regulons, as they partially represented the cartilage lineage module (Figure 2G). On the contrary, we found that MEF2C targets contributed to the chondrocyte differentiation pathway, although these targets had been known for their osteogenic activities (BMP2, BMP4, WNT5B, WNT10B, SOX9, SULF1, RUNX3, FGFR3, RUNX2, RUNX1). This reiterates that MEF2C specifically regulated the early bone cluster, as previously outlined by the composition of its regulon (Figure 2G).

Together, our results indicated that transcriptional programs were distinguished from gene expression. Thus, we capitalized on the advantage of network inference for cell state identification, and coupled with LRLR to assemble novel gene and regulatory signatures of day 0 and day 14 during MSC differentiation.

### Trajectory analysis identified three end-states in MSC differentiation stemming from one starting point

As heterogeneity existed at day 0 and day 14, we set out to further elucidate the differentiation hierarchy of MSCs. We asked whether multiple differentiated states arose from individual committed multiple progenitor pools, or whether these progenitor pools exhibited a level of plasticity, and thus all pools gave rise to all differentiated fates. In the first scenario, specialized progenitor cells are restricted in their multipotency and could only give rise to specific differentiated states (i.e., “nature”). In this case, it would be beneficial to isolate such subpopulations by targeted fluorescence activated cell sorting (FACS) to achieve homogeneity in engineered tissue. Alternatively, in the second scenario, while MSCs in this population exhibit intrinsic differences, molecules from the microenvironment such as culture conditions actually provided the driving force to induce MSCs towards all differentiated lineages (i.e., “nurture”). In this case, dissection of the gene regulatory network to identify decision points during lineage specification is critical, as this method would aid in designing an alternative protocol to induce MSCs towards one specific lineage.

On a single-cell level, the “nature” versus “nurture” path exhibited very distinct profiles, either by cellular projection on a UMAP space or by reconstructed trajectory (Figure 3A). We found that both UMAP projection and trajectory analysis of MSC differentiation pointed towards the nurture scenario, where cells started at one state, and then differentiated into three end-states of preserved stromal progenitors, cartilage/early bone, and hypertrophic cartilage (Figure 3B). The specification to progenitor pool happened early on in differentiation, followed by a later split to either cartilage or hypertrophic cartilage. Next, we inspected the expression of lineage module markers throughout our inferred trajectory (Figure 3C), and showed that the expression of cartilage, early bone and hypertrophic modules corresponded to the cartilage and hypertrophic branches, respectively. Cells previously labeled as “immature” were in fact characterized by a transition state, where multipotent MSCs were differentiating to either bone or hypertrophic cartilage. We also inspected each branch for expression of canonical markers and confirmed that the preserved stromal branch expressed LEPR and LUM (Figure S3) (21, 22).

**Figure 3:**
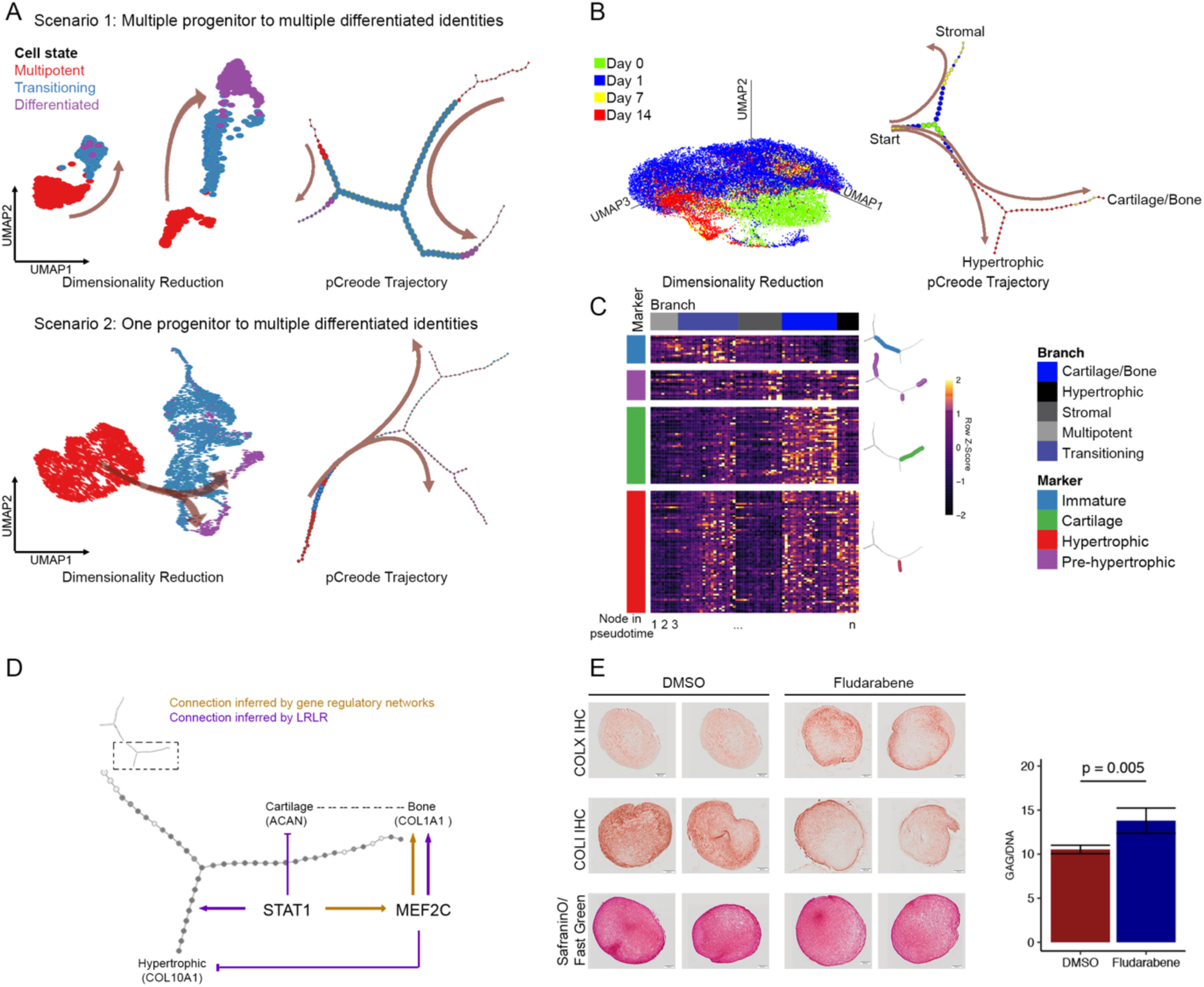
MSC differentiation is a nurture process. (A) Proposed scenarios of MSC chondrogenesis and the respective outcomes with UMAP projection and pCreode trajectory. (B) Both UMAP projection and pCreode trajectory pointed towards a “nurture” scenario instead of “nature” in MSC differentiation. (C) Expression of lineage markers along differentiation branches. (D) STAT1 and MEF2C regulations of cartilage/bone and hypertrophic branches. (E) Phenotypic outcomes of fludarabene treatments on day 14 MSC pellets. n = 5. Welch’s t-test.

Concomitantly, the cartilage branch also expressed COL10A1, and seemed to start expressing COL1A1, confirming that this branch was en-route to becoming hypertrophic, then osteoblastic. The hypertrophic branch, on the other hand, exhibited unique expression of many genes relating to transcription and translation (such as EIF3I and TCEB1) (Figure S3). Recent findings on skeletal system development suggest that hypertrophic chondrocytes could transdifferentiate into osteoblasts or progress towards apoptosis (23, 24). Our cartilage/early bone branch indicated that *in vitro* MSC differentiation may follow the path of chondrocyte-to-osteoblast transdifferentiation. These findings raise the possibility that the hypertrophic branch represents an earlier snapshot of this process, during which chondrocytes show increased protein synthesis. Cells constituting this branch could face the ultimate fate of death, or they may be analogous to the preserved alternative source that could transdifferentiate into osteoblasts upon fracture healing (25).

#### Decision point between osteoblastic and hypertrophic fates was governed by STAT1

Our trajectory analysis indicated that MSC differentiation is a nurture process, and thus proposed a framework where alternative conditions could result in desired tissue phenotypes, by inducing one differentiated branch while constraining the others. To this end, we investigated the inferred LRLR results and gene regulatory networks to identify transcription factor candidates. We reasoned that if a candidate was specific to one branch, its pharmacological inhibition would result in branch elimination, ultimately driving multipotent cells towards other fates. Both STAT1 and IRF3 exhibited inducing potentials for hypertrophic identity (day14 – TFA1 cluster), while displaying inhibitory effects for cartilage identity (day14 – TFA3 cluster) (SI 1). STAT1 was in turn an upstream regulator of MEF2C, a transcription factor with established link to COL1A1 activation (26). Interestingly, MEF2C also possessed negative weight for terminal hypertrophy. Altogether, STAT1 and MEF2C exemplified the transcription factors of interest, portraying a simplistic snapshot of the GRN involved in governing between cartilage/early bone and hypertrophic fates (Figure 3D).

We hypothesized that pharmacological inhibition of STAT1 by fludarabene will result in reduced hypertrophy and enhanced cartilage formation. Subsequently, MEF2C will decrease in the absence of STAT1. This would be expected to inhibit osteogenic differentiation, but also to cause de-repression of the hypertrophic identity. To test our prediction, we induced MSCs towards chondrogenesis in the presence of fludarabene (2.5 μM versus DMSO control), and investigated pellet outcomes at day 14 by histological and biochemical assays. Indeed, there was a decrease in collagen type I staining for pellets cultured in fludarabene, coupled with higher GAG production (Figure 3E). As predicted, we also observed an increase in collagen type X as a side effect, corresponding with de-repression of the hypertrophic branch by MEF2C deficiency.

In the minimal snapshot of STAT1 and MEF2C, we were able to reproduce outcomes predicted by both LRLR and GRN. While the gene regulatory network governing hypertrophy versus cartilage formation is complicated, our results demonstrated a promising approach where network dissection will ultimately result in optimized conditions for cartilage tissue engineering.

### Similarities and variations across individual scRNA-seq data

An important question is whether cellular architecture and differentiation trajectory are similar across MSCs from different individuals. To answer this question, we performed scRNA-seq following MSC differentiation using a different donor source. Since sources were de-identified, information regarding sex and age was unknown. However, expression of senescence markers was similar between the two donors (Figure S4B), and based on the expression of JPX and XIST (Figure S4B), we speculate that donor 1 (reported thus far in this manuscript) was female and donor 2 was male. Similar to donor 1, donor 2 exhibited distinct cellular clustering based on transcription factor activity versus gene expression (Figure S4A): about 3-5 TFA clusters and 3-4 GE clusters were identified per time point.

To infer the cell types contributing to donor 2 day 14, we first visualized cluster structures from two disparate data sets on a shared space (Figure 4A) and detected the presence of all four previously proposed identities. Next, each cell from donor 2 day 14 was assigned a prediction score either by transfer learning (embedded Seurat function (13)) or by LRLR (Figure 4B). Both methods identified TFA1 as the cartilage cluster, TFA2 and TFA5 as the immature, TFA3 as the early bone, and TFA4 as the hypertrophic cluster. We observed a strong agreement between cellular identities inferred by Seurat transfer label and LRLR (81%; 1064 in 1319 cells). Cells with dissimilar labels were subsequently investigated further for lineage module expression.

**Figure 4:**
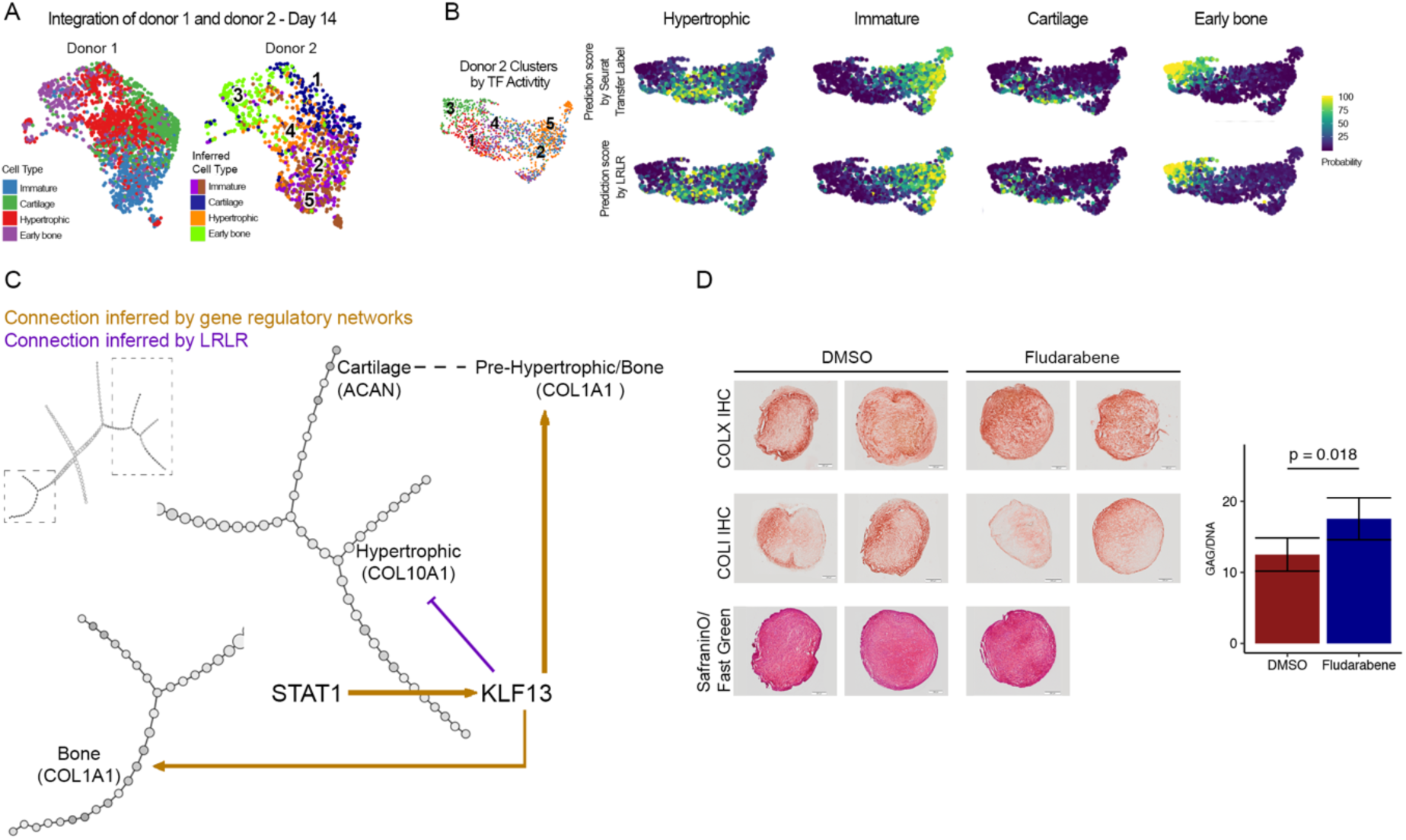
Characteristics of donor 2 MSC differentiation. (A) Integration of donor 1 and donor 2 data set onto a shared space. (B) Cell type prediction with Seurat transfer label and lasso regularized logistic regression (LRLR). (C) STAT1 and KLF13 regulation of cartilage/bone and hypertrophic branches. (D) Phenotypic outcomes of fludarabene treatments on day 14 MSC pellets from donor 2. n = 5. Welch’s t-test.

Interestingly, cells proposed as immature by Seurat transfer label exhibited high levels of unique hypertrophic markers EIF3I and TCEB1, while cells proposed as hypertrophic displayed low expression of both genes. This discrepancy was not observed with cell type inference by LRLR (Figure S4C), and thus indicates that at least in the case of MSC differentiation, LRLR could be more accurate in representing biological insights from one individual data set to the other.

Donor 2 followed a similar trajectory to donor 1, corroborating the nurture scenario during MSC differentiation. We observed an earlier split to preserved progenitor pool, followed by a later split to either cartilage/maturing bone or hypertrophic cartilage (Figure S5). However, there also exists an independent branch forming early on and progressing to bone – a phenomenon not observed in donor 1. This branch was predominantly constituted by donor 2 day 0 TFA5 (Figure S5D), a cluster heavily influenced by SRF transcriptional activity that was directly upstream of many osteoblastic markers such as SULF1 and LRP1 (SI 2) (27–30). We speculate that this particular branch may follow the intramembranous, instead of the endochondral ossification route as observed later on in the trajectory. While the endochondral ossification path demonstrated similarity across donors, the presence of an intramembranous ossification domain underlined individualized distinctions in cellular architecture – an important feature that could have been overshadowed by bulk RNA sequencing.

Finally, we examined whether STAT1 influence was recapitulated in donor 2. STAT1 did not exhibit heavy weights for any differentiated identities, but was directly upstream of KLF13. LRLR suggested an inhibitory effect of KLF13 on the hypertrophic cluster, and inferred GRN pinpointed that KLF13 upregulated COL1A1 (Figure 4C). Taken together, it appears that STAT1 may act through KLF13 to enhance the bone identity, and thus inhibition of STAT1 by fludarabene could result in decreased bone outcome. Collaterally, loss of KLF13 activity would lead to de-repression of hypertrophic identity, and thus increased hypertrophic outcome. While no connection was directly drawn to the cartilage cluster, cartilage outcome could potentially improve upon inhibition of the other paths. Indeed, fludarabene-treated pellets at day 14 displayed increased GAG production and decreased COLI staining, while COLX staining may be similar or slightly increased compared to the DMSO control (Figure 4D). While certain levels of donor-to-donor variation exist, our results demonstrated that STAT1 held a crucial function in MSC differentiation in a donor-independent manner.

## Discussion

We used scRNA-seq to determine the transcriptomic and regulatory landscapes of MSC differentiation at the single cell level. Cellular heterogeneity was observed at all time points in our analysis and was better represented by distinctive transcriptional programs rather than by variations in gene expression. We demonstrated the use of LRLR in (1) identifying gene as well as regulatory signatures for each cluster (immature, cartilage, hypertrophic cartilage, and early bone), and (2) classifying cell identities. LRLR-derived lineage module markers were more distinguished between clusters compared to those identified by Wilcoxon rank sum test (Seurat FindMarkers default method). In the case of MSC differentiation, classification of cell identities by LRLR also offered us more relevant biological insights compared to Seurat transfer label.

MSC heterogeneity is an increasingly appreciated subject. In fact, primary, passage-2 MSCs-derived clonal sub-populations had been shown to exhibit tri-lineage, bi-lineage, or uni-lineage potentials (31). However, as MSCs undergo population doublings during culture expansion, their proliferation decreases and so does their multipotency (31–34). Indeed, functional variation in later passages is represented by a dwindling number of multipotent stem cells and the emergence of an osteochondral progenitor subpopulation (32, 33). As MSCs of later passage were utilized in our study (passage 6), we speculate that supplemented molecules in culture conditions had instructed multipotent MSCs towards both osteogenic and chondrogenic lineages, but restricted their differentiation towards other pathways. Moreover, inherent, existing osteochondral progenitor subpopulation may also thrive in this defined environment. Consequently, the heterogeneity in our day 0 population can be explained by a potency hierarchy, where cell clusters represent MSCs along the path of multipotent to osteochondral progenitor restriction. We also speculate that other non-chondrogenic clones (adipogenic or osteo-adipogenic) may be out-competed by osteochondral progenitors over passages or during pellet culture, and thus did not constitute a significant cluster in our analysis.

Inferred trajectory revealed that MSC differentiation is not a process where progenitors displayed restricted ability to differentiate into certain lineages, but rather a process where exogenous factors instructed multipotent cells towards various end-states. Interestingly, trajectory of MSC differentiation may vary from donor to donor, as our results indicated a phase of intramembranous ossification in donor 2 but not donor 1. Nevertheless, influential transcriptional programs were comparable across individuals, as in the case of STAT1. Here, we showed that STAT1 acted through MEF2C in donor 1 and KLF13 in donor 2 to enhance the bone identity. It is worth noting that connection from STAT1 to both MEF2C and KLF13 could exist in both donors and may be lost with the current computational method used to define regulons (8). While the roles of both MEF2C in skeletal development had been extensively studied, KLF13 and STAT1 function remained poorly understood.

In summary, the LRLR approach provides several advances in the analysis of scRNA-seq data by identifying of small number of highly influential genes or transcription factors, as well as providing a means of classifying similar but distinct cell types within a population. Our results provide a proof of principal that single cell RNA sequencing combined with LRLR and GRN reconstruction could identify novel targets for either pharmacological intervention or gene perturbation to enhance the quality of engineered tissue constructs.

## Experimental Methods

### MSC chondrogenesis and drug treatment

Mesenchymal stem cell collection and chondrogenic induction was carried out as previously described (7). In short, bone marrow aspirates from de-identified donors were collected with approval of the Institutional Review Board of Duke University Medical Center. Expansion medium was composed of DMEM-low glucose (Thermo Fisher Scientific), 1% penicillin/streptomycin (Thermo Fisher Scientific), 10% lot-selected fetal bovine serum (FBS; Thermo Fisher Scientific), and 1 ng/ml basic fibroblast growth factor (SigmaAldrich).

Chondrogenic medium was composed of DMEM-high glucose (Thermo Fisher Scientific), 1% penicillin/streptomycin (Thermo Fisher Scientific), 1% Insulin-Transferrin-Selenium Plus Premix (ITS+) (Corning, Corning, NY, USA), 100 nM dexamethasone (Sigma-Aldrich), 50 mg/ml ascorbic acid (Sigma-Aldrich), 40 mg/ml L-proline (Sigma-Aldrich), and 10 ng/ml recombinant human transforming growth factor beta 3 (rhTGF-b3) (R&D Systems). Pellets were formed at the end of passage 6. For drug treatment experiments, pellets were cultured in chondrogenic medium supplemented with either DMSO or 2.5 μM fludarabene. Pellets were assessed at day 14 with biochemical and histological assays as previously described (7).

### Cell isolation and scRNA-seq

On the day of harvest, pellets were digested in 750 μl of collagenase/pronase solution at 37ºC for up to 45 minutes in 15-minute increments, with gentle agitation. Digestion solution was composed of 780 U/ml pronase (EMD Millipore), 1220 U/ml collagenase (Type II, Worthington-Biochem), 5% FBS in DMEM-HG. After matrix degradation, cells were collected and centrifuged at 200 × g for 6 minutes at room temperature. Subsequently, cells were rinsed in 1 ml of PBS and centrifuged again at 200 × g for 5 minutes. Matrix and debris usually collected at the top edge of the tubes, and thus were eliminated by aspiration. Rinsing was repeated one more time with 400 μl of PBS. At this point, cells were counted and resuspended at 2 × 10^6^ cells/ml for cryopreservation in freeze medium (80% FBS, 10% DMEM-HG, 10% DMSO).

On the day of capture, cells were recovered from freeze medium by resuspension in 10ml of 10% FBS/ DMEM-HG. The cell mixture was centrifuged at 200 × g for 5 minutes and rinsed in 1 ml of 0.04% bovine serum albumin/PBS. Cells were again centrifuged at 150 × g for 3 minutes. The rinsing step was repeated one more time. Finally, cells were resuspended at 1,000 cells/µl in 0.04% bovine serum albumin/PBS and ready for microfluidic capture.

Microfluidic capture on the 10× Chromium Controller (10× Genomics), subsequent library preparation using Chromium Single Cell 3’ v2 Reagent kit, and sequencing on the Illumina NovaSeq S1 platform (Illumina) were performed by the Genome Technology Access Center at Washington University in St Louis.

### scRNA-seq analysis

#### Read alignment

Raw sequencing was processed and aligned to the human genome assembly (hg19) using Cell Ranger software (v2, 10x Genomics).

#### Preprocessing steps

## Filtering

Cells with high mitochondrial content (5% of the total reads) were removed (Seurat 2.3.4) (35). Additionally, cells with low RNA recovery or very high RNA content (doublets) were also excluded from downstream analysis (Monocle 2.6.0) (36).

## Normalization and data scaling

Technical variations such as sequencing depth, proportion of mitochondrial transcripts, different phases in non-dividing cell cycles (variation in dividing versus non-dividing cells was retained) were regressed out during data normalization and scaling following instructions from the Seurat package.

## Estimation of cluster numbers

Number of clusters was determined using SIMLR_Estimate_Number_of_Clusters (SIMLR 1.4.1) (37) on normalized gene expression data set or normalized transcription factor activity AUC matrix.

## Determination of significant principal component

Jackstraw function (Seurat 2.3.4) was utilized to indicate significant number of principal components for downstream analysis (p < 0.0001).

## Cell clustering and dimensionality reduction

For gene expression data, FindClusters (Seurat 2.3.4) was performed with the above pre-determined number of clusters and number of principal components.

For transcription factor activity data, SIMLR_Large_Scale (SIMLR 1.4.1) was performed with the above pre-determined number of clusters. Number of principal components were determined by the elbow plot method.

Dimensionality reduction was performed on gene expression data with Uniform Manifold Approximation and Projection (UMAP) (Seurat 2.3.4; umap-learn 0.3.7) (38).

### Determining cluster markers with Seurat

Cluster markers were determined using FindAllMarkers (Seurat 3.0.0) (adjusted p-val < 0.05). The Wilcoxon rank sum test was the default test parameter used for this function.

### Integration of single cell data sets and label transfer

Cell type inference with integration of data sets or label transfer was carried out following instruction from the Seurat package (Seurat 3.0.0).

### Inference of gene regulatory networks

Transcription factors and related regulons were computed with pySCENIC following package instruction (8). Pre-computed reference provided with pySCENIC includes database ranking and motif annotation for *Homo sapiens*.

#### Trajectory analysis

Equal number of cells from each time point was selected randomly, and combined to create balanced data. Additionally, data was further subset to contain the combination of all lineage module markers as features. This was the input for pCreode, and the proposed trajectory was built following the package instructions (39). Output graphs were scored and the highest (i.e. most representative) graphs were reported in this manuscript.

### Lasso regularized logistic regression

We used lasso regularized logistic regression (LRLR) with one-versus-all strategy from the package lassoglm to classify cells from different cell types. For each cell type, the response variable was coded as 1 if a cell belonged to that cell type and 0 otherwise. The optimal tuning parameter was chosen via ten-fold cross-validation. The resulting models were sparse with weights for genes in the raw count matrix. For a new cell expression data, the models were used to calculate the probability that the cell belonged to a particular cell type. The new cell was assigned to the cell type with highest probability. From our cross-validation study, LRLR achieved average accuracy of 90% and average AUC of 0.96.

#### Statistical analysis

Statistical analysis for biochemical assay outcomes was performed using R (Vienna, Austria).

## Acknowledgments

Supported in part by grants from the NIH (AR50245, AR46927, AG15768, AR48182, AR057235, AR073752), Arthritis Foundation, and Nancy Taylor Foundation for Chronic Diseases. We thank the Genome Technology Access Center in the Department of Genetics at Washington University School of Medicine. The authors thank Dr. Samantha Morris for discussions in the early stages of this work, and Sara Oswald for assistance with scientific writing.

## Author contribution

NPTH, FG designed the study. NPTH, NHK conducted the study and collected data for single cell RNA sequencing. DBK conducted study and collected data for drug treatment validation. NPTH, MP analyzed data. NPTH wrote the manuscript. All authors edited manuscript content.

---

**Figure S1:**
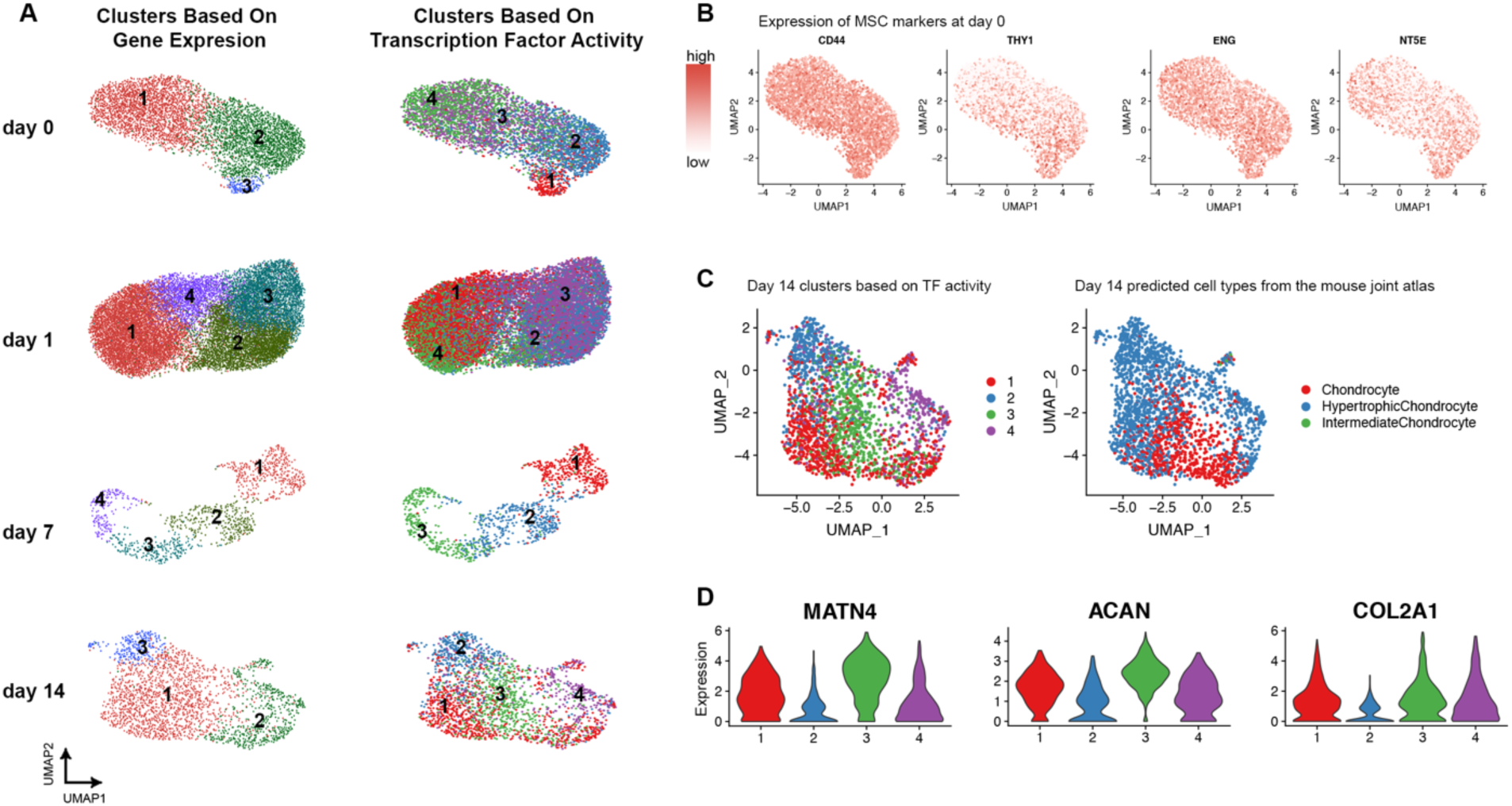
Overview of MSC differentiation scRNA-seq data set. (A) Cellular clustering based on gene expression or transcription factor activity for four investigated time points. (B) MSC markers’ expression at day 0. (C) Cell type prediction of day 14 clusters using Seurat label transfer with the mouse joint atlas as reference. (D) Expression of canonical cartilage markers in day 14 clusters.

**Figure S2:**
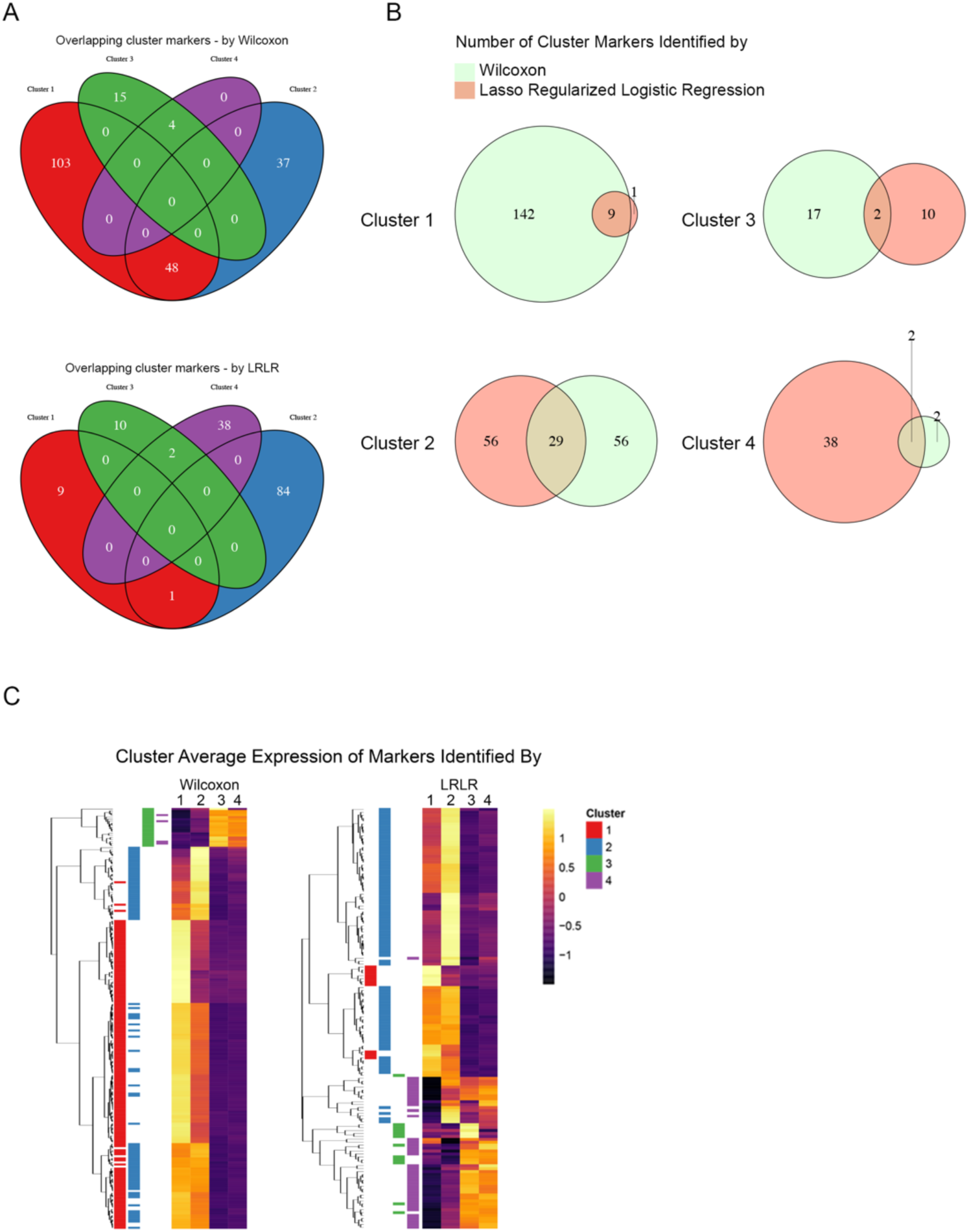
Wilcoxon rank sum test versus lasso regularized logistic regression to identify markers for each cell cluster in day 0. (A) Venn diagram showing number of overlapping cluster markers by each test. (B) Venn diagram showing number of overlapping markers between the two tests. (C-D) Heatmap depicting gene signatures for each cluster, color-mapped by gene average expression per cluster. Each heatmap row represents a gene, and is annotated with the cluster of which gene was designated as marker.

**Figure S3:**
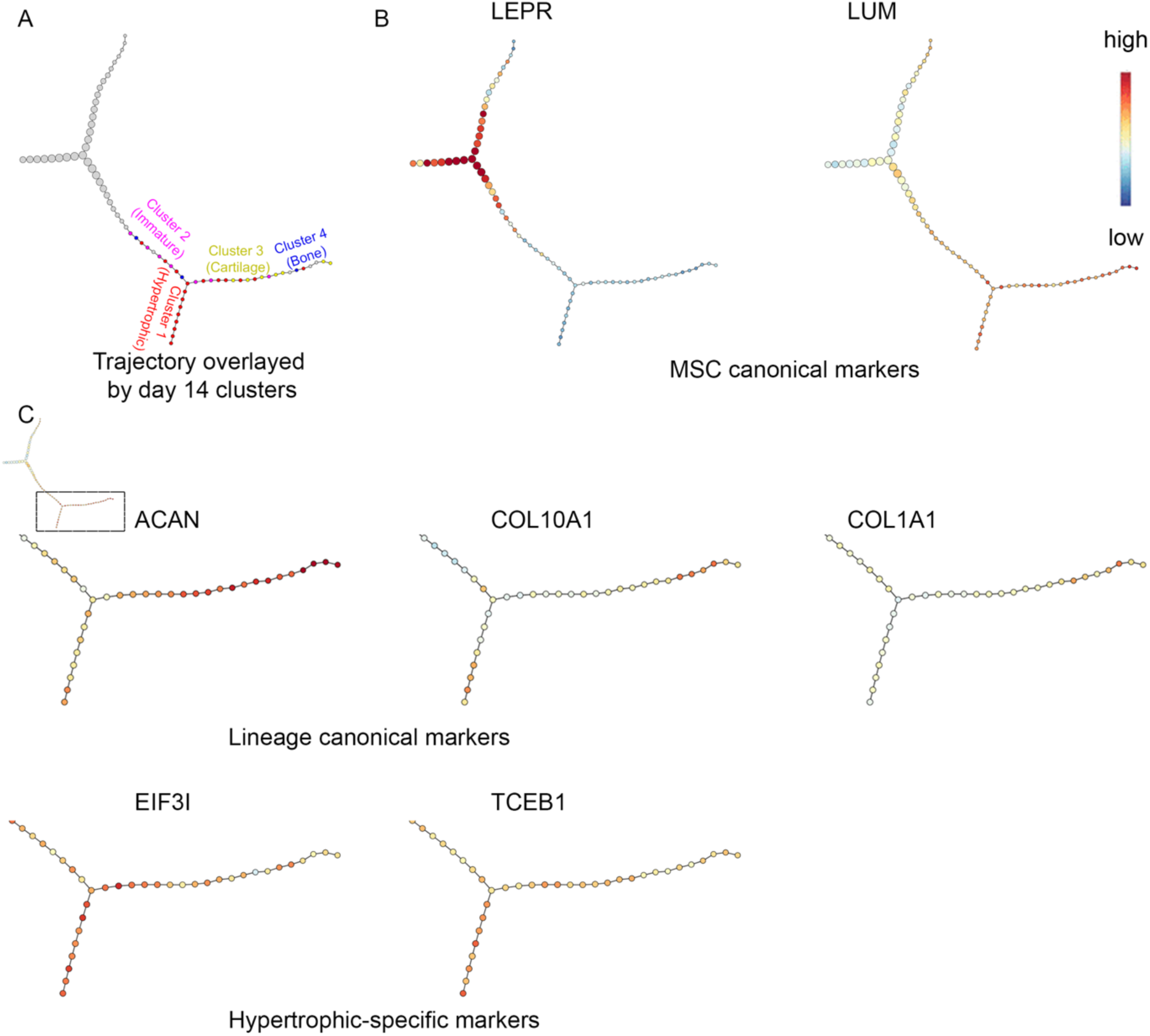
Proposed trajectory of MSC differentiation in donor 1. (A) Position of day 14 clusters on the trajectory. (B) Expression of canonical MSC markers throughout trajectory. Expression level is scaled within each gene. (C) Expression of canonical lineage markers and hypertrophic specific markers.

**Figure S4:**
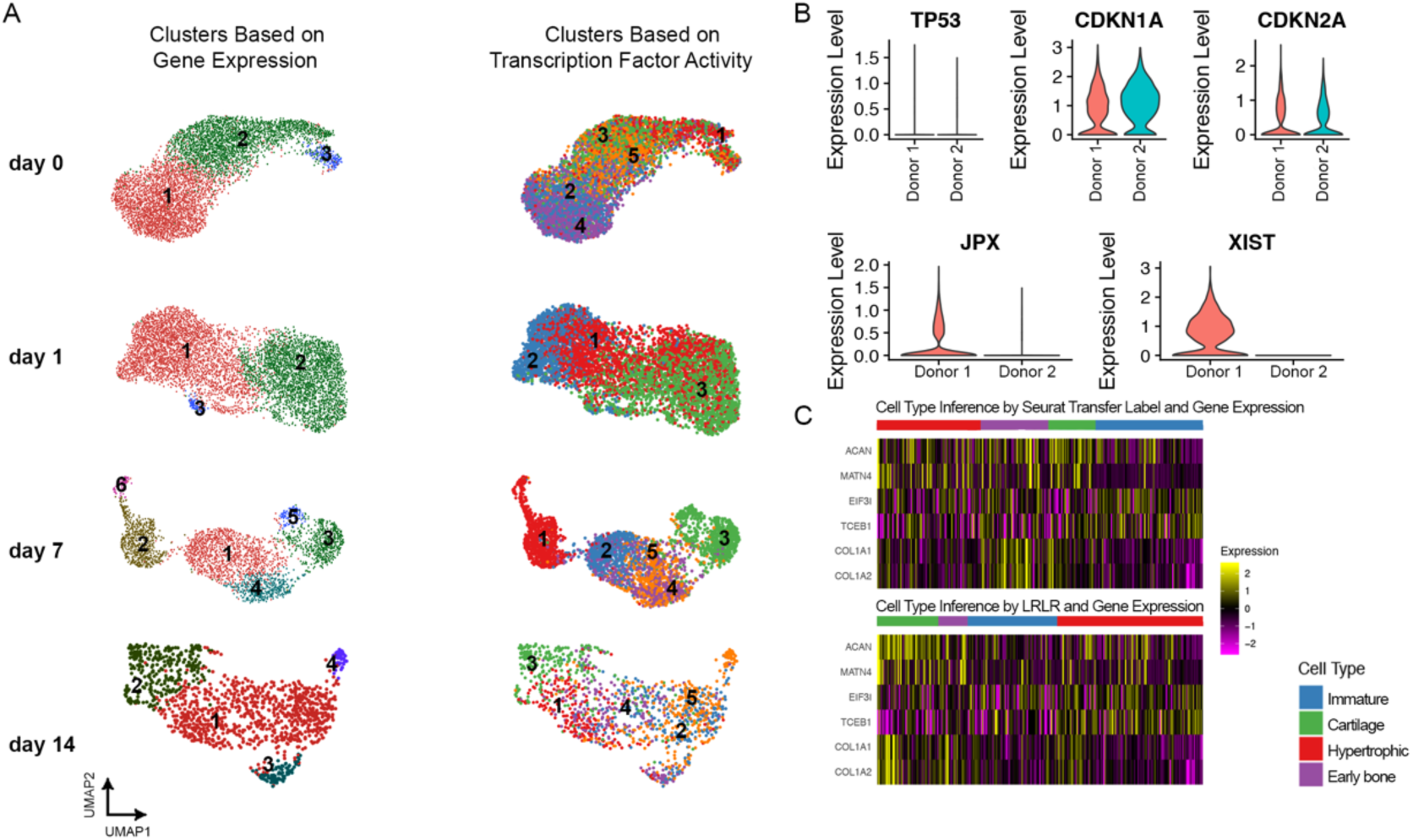
Overview of MSC differentiation scRNA-seq data set from donor 2. (A) Cellular clustering based on gene expression or transcription factor activity for four investigated time points. (B) Expression of senescence and sex-related genes in donor 1 versus donor 2. (C) Expression of lineage markers and inferred cell type classification by Seurat transfer label or by LRLR.

**Figure S5:**
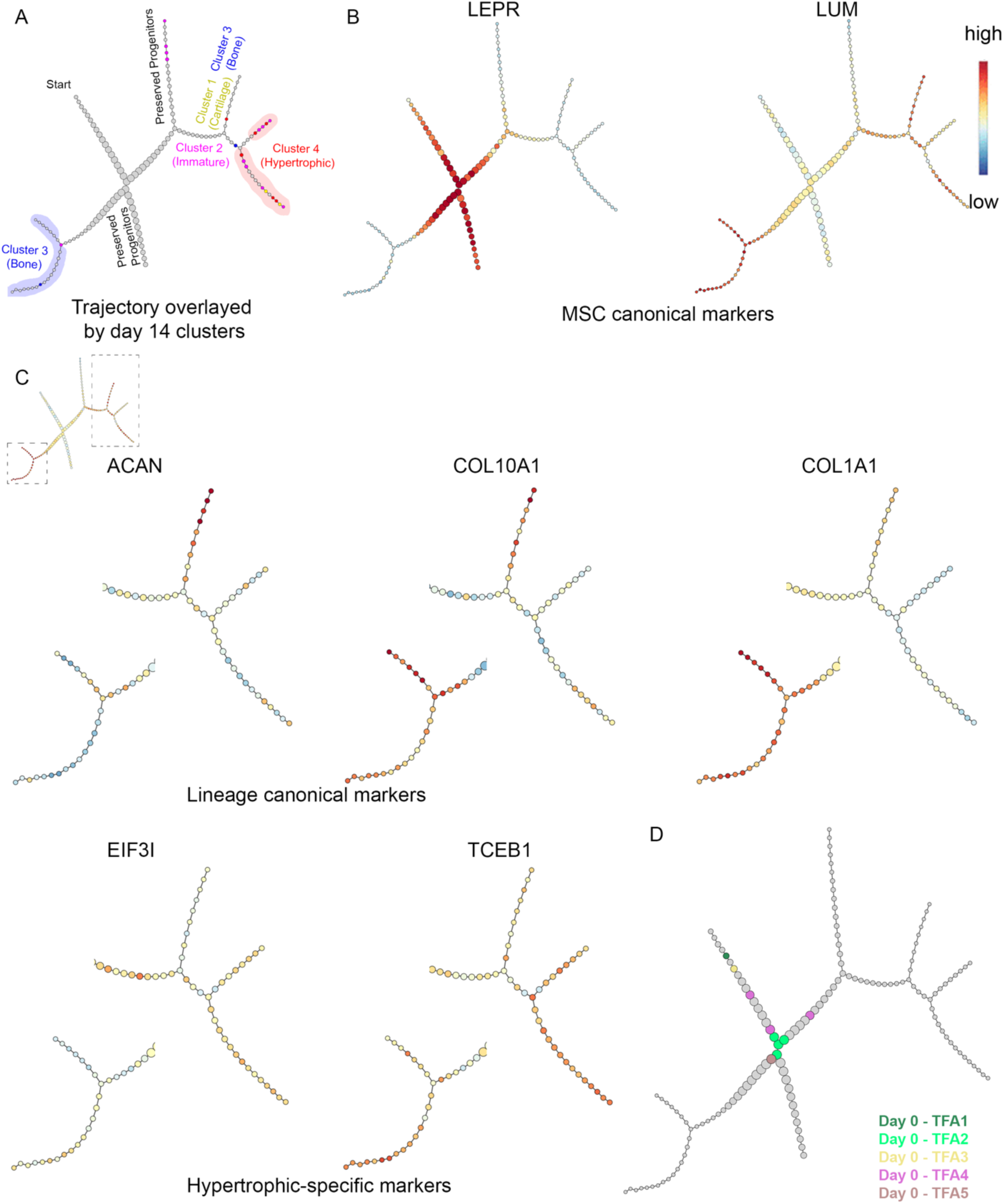
Proposed trajectory of MSC differentiation in donor 2. (A) Position of day 14 clusters on the trajectory. (B-C) Expression of canonical MSC markers, lineage markers, and hypertrophic specific markers. Expression level is scaled within each gene. (D) Position of day 0 clusters on the trajectory.

## References

1. Heinemeier KM, et al. (2016) Radiocarbon dating reveals minimal collagen turnover in both healthy and osteoarthritic human cartilage. Sci Transl Med 8(346):346ra390.

2. Nguyen US, et al. (2011) Increasing prevalence of knee pain and symptomatic knee osteoarthritis: survey and cohort data. Ann Intern Med 155(11):725–732.

3. Johnstone B, Hering TM, Caplan AI, Goldberg VM, & Yoo JU (1998) In vitro chondrogenesis of bone marrow-derived mesenchymal progenitor cells. Exp Cell Res 238(1):265–272.

4. Mackay AM, et al. (1998) Chondrogenic differentiation of cultured human mesenchymal stem cells from marrow. Tissue Eng 4(4):415–428.

5. Mueller MB, et al. (2010) Hypertrophy in mesenchymal stem cell chondrogenesis: effect of TGF-beta isoforms and chondrogenic conditioning. Cells Tissues Organs 192(3):158–166.

6. Mueller MB & Tuan RS (2008) Functional characterization of hypertrophy in chondrogenesis of human mesenchymal stem cells. Arthritis Rheum 58(5):1377–1388.

7. Huynh NPT, Zhang B, & Guilak F (2019) High-depth transcriptomic profiling reveals the temporal gene signature of human mesenchymal stem cells during chondrogenesis. FASEB J 33(1):358–372.

8. Aibar S, et al. (2017) SCENIC: single-cell regulatory network inference and clustering. Nat Methods 14(11):1083–1086.

9. Houle M, Prinos P, Iulianella A, Bouchard N, & Lohnes D (2000) Retinoic acid regulation of Cdx1: an indirect mechanism for retinoids and vertebral specification. Mol Cell Biol 20(17):6579–6586.

10. Subramanian V, Meyer BI, & Gruss P (1995) Disruption of the murine homeobox gene Cdx1 affects axial skeletal identities by altering the mesodermal expression domains of Hox genes. Cell 83(4):641–653.

11. Inada M, et al. (2004) Critical roles for collagenase-3 (Mmp13) in development of growth plate cartilage and in endochondral ossification. Proc Natl Acad Sci U S A 101(49):17192–17197.

12. Dy P, et al. (2008) The three SoxC proteins–Sox4, Sox11 and Sox12–exhibit overlapping expression patterns and molecular properties. Nucleic Acids Res 36(9):3101–3117.

13. Stuart T, et al. (2019) Comprehensive Integration of Single-Cell Data. Cell 177(7):1888–1902 e1821.

14. Kelly NH, Huynh, N.P.T., Guilak, F. (2019) Single cell RNA-sequencing reveals cellular heterogeneity and trajectories of lineage specification during murine embryonic limb development. bioRxiv.

15. Florine EM, et al. (2013) Effects of Dexamethasone on Mesenchymal Stromal Cell Chondrogenesis and Aggrecanase Activity: Comparison of Agarose and Self-Assembling Peptide Scaffolds. Cartilage 4(1):63–74.

16. Derfoul A, Perkins GL, Hall DJ, & Tuan RS (2006) Glucocorticoids promote chondrogenic differentiation of adult human mesenchymal stem cells by enhancing expression of cartilage extracellular matrix genes. Stem Cells 24(6):1487–1495.

17. Ikeda T, et al. (2004) The combination of SOX5, SOX6, and SOX9 (the SOX trio) provides signals sufficient for induction of permanent cartilage. Arthritis Rheum 50(11):3561–3573.

18. Lefebvre V, Huang W, Harley VR, Goodfellow PN, & de Crombrugghe B (1997) SOX9 is a potent activator of the chondrocyte-specific enhancer of the pro alpha1(II) collagen gene. Mol Cell Biol 17(4):2336–2346.

19. Lefebvre V, Li P, & de Crombrugghe B (1998) A new long form of Sox5 (L-Sox5), Sox6 and Sox9 are coexpressed in chondrogenesis and cooperatively activate the type II collagen gene. EMBO J 17(19):5718–5733.

20. Sitara D & Aliprantis AO (2010) Transcriptional regulation of bone and joint remodeling by NFAT. Immunol Rev 233(1):286–300.

21. Raouf A, et al. (2002) Lumican is a major proteoglycan component of the bone matrix. Matrix Biol 21(4):361–367.

22. Zhou BO, Yue R, Murphy MM, Peyer JG, & Morrison SJ (2014) Leptin-receptor-expressing mesenchymal stromal cells represent the main source of bone formed by adult bone marrow. Cell Stem Cell 15(2):154–168.

23. Yang L, Tsang KY, Tang HC, Chan D, & Cheah KS (2014) Hypertrophic chondrocytes can become osteoblasts and osteocytes in endochondral bone formation. Proc Natl Acad Sci U S A 111(33):12097–12102.

24. Shapiro IM, Adams CS, Freeman T, & Srinivas V (2005) Fate of the hypertrophic chondrocyte: microenvironmental perspectives on apoptosis and survival in the epiphyseal growth plate. Birth Defects Res C Embryo Today 75(4):330–339.

25. Aghajanian P & Mohan S (2018) The art of building bone: emerging role of chondrocyte-to-osteoblast transdifferentiation in endochondral ossification. Bone Res 6:19.

26. Arnold MA, et al. (2007) MEF2C transcription factor controls chondrocyte hypertrophy and bone development. Dev Cell 12(3):377–389.

27. Chen J, et al. (2012) Serum response factor regulates bone formation via IGF-1 and Runx2 signals. J Bone Miner Res 27(8):1659–1668.

28. Liu TM & Lee EH (2013) Transcriptional regulatory cascades in Runx2-dependent bone development. Tissue Eng Part B Rev 19(3):254–263.

29. Holst CR, et al. (2007) Secreted sulfatases Sulf1 and Sulf2 have overlapping yet essential roles in mouse neonatal survival. PLoS One 2(6):e575.

30. Niemeier A, et al. (2005) Expression of LRP1 by human osteoblasts: a mechanism for the delivery of lipoproteins and vitamin K1 to bone. J Bone Miner Res 20(2):283–293.

31. Russell KC, et al. (2010) In vitro high-capacity assay to quantify the clonal heterogeneity in trilineage potential of mesenchymal stem cells reveals a complex hierarchy of lineage commitment. Stem Cells 28(4):788–798.

32. Lee WC, et al. (2014) Multivariate biophysical markers predictive of mesenchymal stromal cell multipotency. Proc Natl Acad Sci U S A 111(42):E4409–4418.

33. Whitfield MJ, Lee WC, & Van Vliet KJ (2013) Onset of heterogeneity in culture-expanded bone marrow stromal cells. Stem Cell Res 11(3):1365–1377.

34. Rennerfeldt DA & Van Vliet KJ (2016) Concise Review: When Colonies Are Not Clones: Evidence and Implications of Intracolony Heterogeneity in Mesenchymal Stem Cells. Stem Cells 34(5):1135–1141.

35. Butler A, Hoffman P, Smibert P, Papalexi E, & Satija R (2018) Integrating single-cell transcriptomic data across different conditions, technologies, and species. Nat Biotechnol 36(5):411–420.

36. Trapnell C, et al. (2014) The dynamics and regulators of cell fate decisions are revealed by pseudotemporal ordering of single cells. Nat Biotechnol 32(4):381–386.

37. Wang B, et al. (2018) SIMLR: A Tool for Large-Scale Genomic Analyses by Multi-Kernel Learning. Proteomics 18(2).

38. Becht E, et al. (2018) Dimensionality reduction for visualizing single-cell data using UMAP. Nat Biotechnol.

39. Herring CA, et al. (2018) Unsupervised Trajectory Analysis of Single-Cell RNA-Seq and Imaging Data Reveals Alternative Tuft Cell Origins in the Gut. Cell Syst 6(1):37–51 e39.

